# The Potential of Lipolytic Filamentous Fungi Isolated from Landfill Soil as Poly-β-Hydroxybutirate (PHB) Bioplastic Degrader

**DOI:** 10.1101/2019.12.19.883538

**Authors:** Nur Arfa Yanti, Endang Sriwahyuni, Nur Rayani La Omi, Nurhayani H. Muhiddin, Sitti Wirdhana Ahmad

## Abstract

The present study was investigated the potential of lipolytic fungi (molds) isolated from landfill soil in degrading Poly-β-hydroxybutyrate (PHB). Screening of PHB-degrading lipolytic molds was done in two stages, such as screening of lipolytic molds which was identified by the formation of orange fluorescent halos around the colony on rhodamine B agar medium and the degradation PHB ability test was identified by the formation of clear zone around colony on PHB emulsion medium. Characterization of isolates was done based on phenotypic characters and the identification was done by numerical-phenetic analysis. Three lipolytic mold isolates that have ability in degrading PHB bioplastic i.e isolate KC1, KE1 and KE6. These molds have asexual spore form conidia, foot cell, septate hyphae, unbranched conidiophore, and spore mass located at the apex of phialid. The identification results showed that isolate KC1 is identic to *Aspergillus terreus*, KE1 is identic to *Aspergillus niger* and KE6 is identic to *Aspergillus fumigatus*.

## Introduction

Generally, the commercial plastics are made of undegradable materials such as crude oils. They are difficult to be degrade, consequently they will be accumulated continuously over the time. This condition will give negative impact as pollutions in the environment. One of the solutions to anticipate it is the utilization of biodegradable plastic or bioplastic which is eco-friendly, such as poly-β-hydroxybutyrate (PHB) (Bharti and Swetha, 2016). PHB as a member of poly-β-hydroxyalkanoate (PHA) is widely produced at industrial scale.

PHB bioplastic has similar character with the polypropilene synthetic plastic, but PHB bioplastic can be produced using renewable resourches which are easier to be degrade in the environment. In additon, PHB is also biocompatible with alive tissues, so it can be applied in biomedical sector (Rathbone *et al*., 2009). Some studies has been reported that PHB is made of natural substrates by the action of microorganisms (Ramadas *et al*., 2009; Full *et al*., 2006; Lafferty *et al*., 1988). Yanti *et al.* (2013; 2016) has been proposed the utilization of local sago starch from Southeast Sulawesi, Indonesia to produce PHB by bacteria, but its biodegradability is not been known.

PHB bioplastic can be degraded enzimatically by lipases becauce its monomer contains of ester bond. Lipases can catalyze the hydrolysis of this ester bond (Leja and Lewandowicz, 2010). However, the lipases enzyme are very expensive and this becomes an obstacle if they are used. Therefore, an alternative way is needed to solve this, such as exploring microorganisms with the capability to synthesis lipases.

Microorganism which is preferred as a great lipase-producing is molds (Falony *et al*., 2006; Thakur, 2012). There were many studies about the exploration of lipolytic molds, but their potential as PHB-degrading molds is still not well known. One of the potential source to find lipolytic molds is in landfill soil because this place is accumulated by the mixture of many wastes containing oil. The present study was conducted to isolate lipolytic molds from landfill soil, to test the potential of these isolates in degrading PHB and to identify them.

## Materials and Methods

1. Soil sampling Soil sample was obtained from Puuwatu landfill area, Kendari, Southeast Sulawesi, Indonesia. Sampling was collected from 5 zones (A, B, C, D and E) with 3 sampling points each area at a depth ± 20 cm from the ground surface. Zones A, B, C and D are former waste disposal areas and have changed functions while zone E is an active waste disposal area. The soil samples were collected in sterile plastics. Soil sample from 3 sampling points of each zone were composited before they were used.
2. Isolation of lipolytic molds *Potato Dextrose Agar* (PDA) (Merck) medium was used in this isolation step. Ten grams of soil sample was suspended in 90 ml of sterile distilled water and was shaken for 15 minutes. Serial dilution were made until 10^−5^. Then, 100 μl dillutions were spread on medium and were incubated at 30°C for 1-2 weeks. Mold colonies grown were purified on PDA plates and were incubated at 30°C for 7 days. Single colony was inoculated on PDA slants and stored at 4°C.
3. Screening of lipolytic molds Screening lipolytic molds was done qualitatively by growing each isolate on rhodamine-B agar medium and was incubated at 37°C for 7 days (Panuthai *et al.*, 2012). Lipolytic activity was detected by irradiating plates with UV light at 365 nm (Murray, 2010).
4. Biodegradation test of PHB biofilm PHB biofilm was obtained from previous study (Yanti *et al*., 2016). Isolates which have maximal lipolytic activity were grown on plastic emulsion agar medium to determine their biodegradation potential (Nathania and Kuswytasari, 2013). Isolates were incubated on medium using point technique and were incubated 30°C for 2 days. Biodegradation activity was determined by the the formation of clear zone around colony. The ratio of clear zone formed was calculated using the equation bellow (Nishida and Tokiwa, 1993).

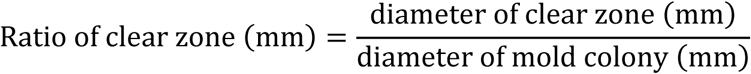
5. Morphology characterization of PHB-degrading lipolytic mold The morphology characterization was carried out macroscopically and microscopically. Macroscopic characteristics observed were the colour of obverse and reverse colony, topography, texture, rate of growth, exudate drops, radial line and diameter of colony. Microscopic characteristics observed were asexual spores, foot cell, septate or aseptate hyphae, conidia, vesicles type and conidiophore characteristics. Microscopic observation was used slide culture method (Sanjaya, 2010; Cappucino and Sharman, 1987). Obtained data were then compared with the descriptions of fungi species in the Training course for the identification of Aspergillus, Penicillium and Talaromyces (Samson, 2017).
6. Identification of PHB-degrading lipolytic mold The identification was done by numerical-fenetic method. The mold isolates and reference species characteristics were analyzed using MVSP (*Multi Variate Statistical Package*) ver.3.1. The diversity of mold fenotipic characteristics were determined by *Simple Matching Coefficient* (SSM) value and the classification were based on UPGMA (*Unweighted Pair Goup Method with Arithmatic Averages)* algoritm. The results were visualized as dendrogram. The dendrogram was used to know the similarity of PHB-degrading lipolytic mold with 4 reference mold species, such as *Aspergillus terreus, Aspergillus niger, Aspergillus flavus* dan *Aspergillus fumigatus.*

## Results

### a. Lipolytic Molds Isolated from Kendari Landfill soil

Twenty four isolates were obtained from Kendari landfill soil. Isolates were screened in rhodamine-B agar medium. Lipolytic activity was indicated as orange fluorescent around the colony by irradiating each isolate in UV light at λ = 365 nm. The higher of the fluorescense intencity, the higher lipoytic activity (Murray, 2010). Sixteen among twenty four isolates were lipolytic molds (Table 1). The difference fluoresent intencity was shown in Figure 1. Three selected lypolytic molds with the highest lipolytic activity were KC1, KE1 and KE6. Hence these isolates were used in the next steps of the study.

**Table 1.**
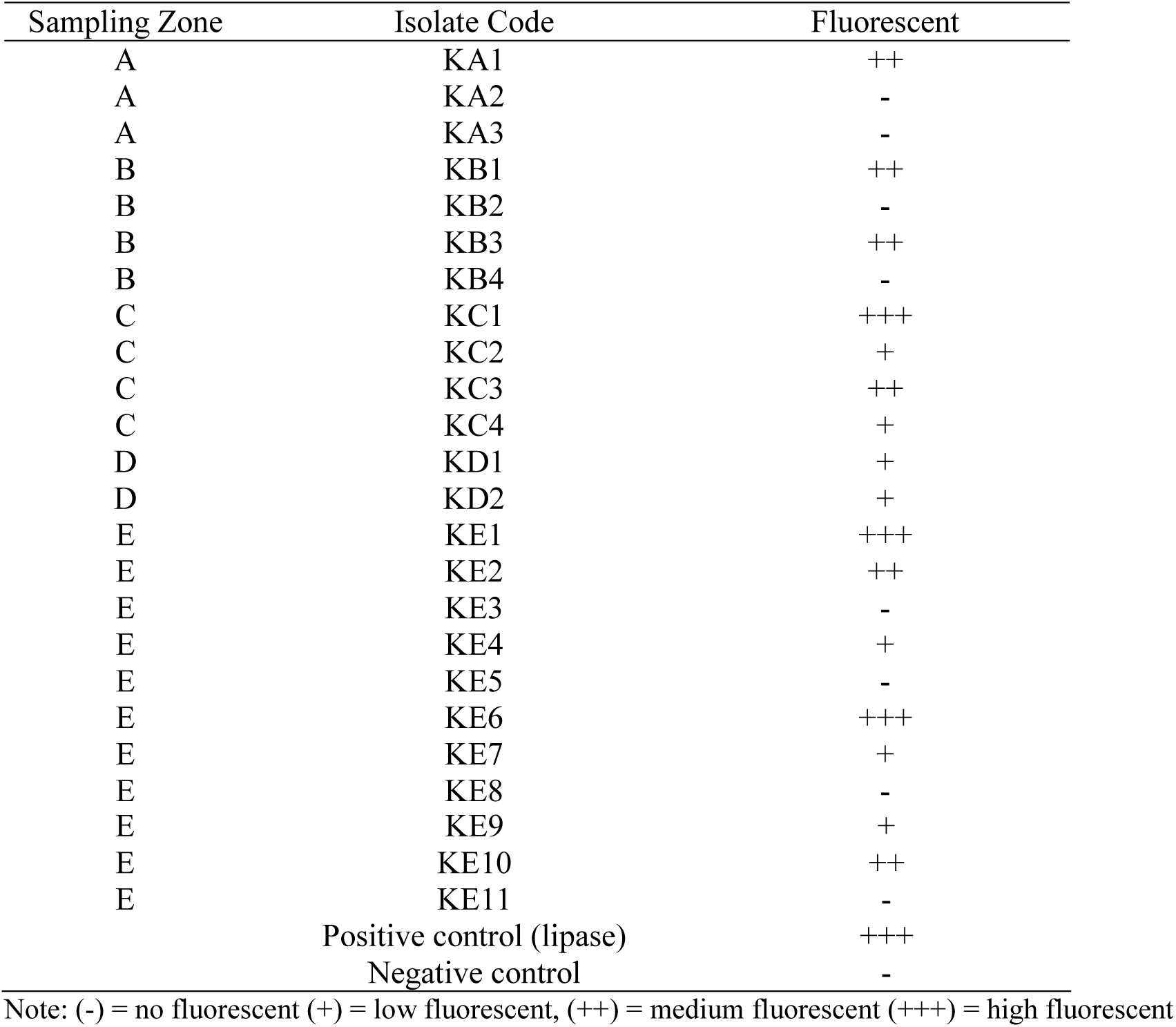
Lipolytic activity based on fluorescent intencity.

**Figure 1.**
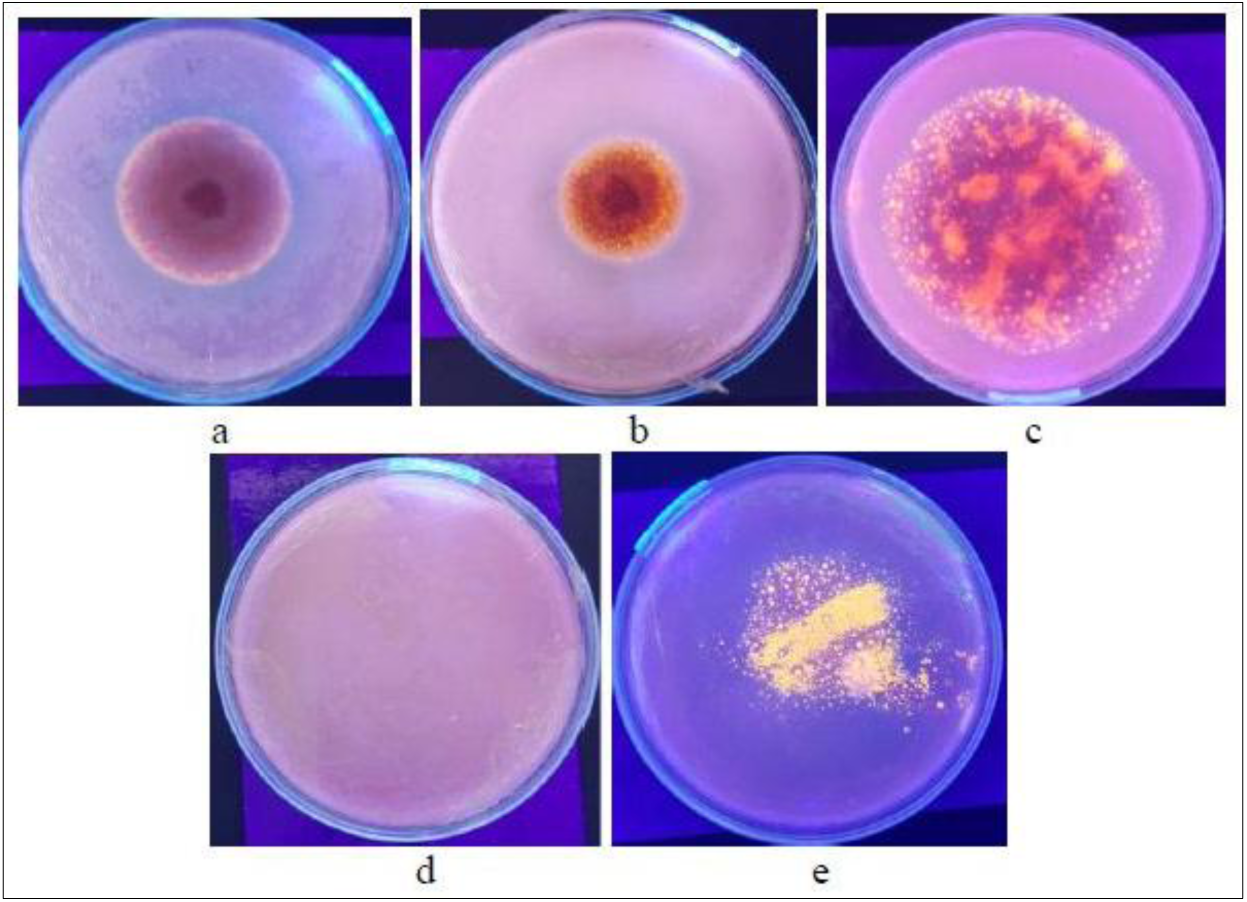
Fluorescent intencity of mold colonies on rhodamine-B agar media, a) low (+), b) medium (++), c) high (+++), d) negative control, e) positive control

### b. Degradation PHB by the selected lipolytic molds

Three selected molds with the highest lipolytic activity were tested in bioplastic emulsion medium to determine their ability in degrading poli-β-hidroksibutirat (PHB). Bioplastic film (PHB) used in this study was obtained from previous study by Yanti *et al*. (2009). Clear zone around the colony indicates that the isolate can degrade poly-β-hidroksibutirate (PHB) (Figure 2). The result of clear zone ratio measurement was shown in Table 2. According to the result obtained, all isolates had the ability to degrade PHB, but isolate KE6 was significantly different to the isolate KC1 and KE6, while isolate KC1 and KE6 were not significantly different at 95% confidence level (Table 2). Three degrading PHB lipolytic molds were further characterized and indentified to determine their identities.

**Table 2.** Degradation ability of three lipolytic molds after 2 days.

| Isolate Code | Diameter of clear zone (mm) | Diameter of colony (mm) | Clear zone ratio |
| --- | --- | --- | --- |
| KC1 | 15 | 12 | 1.25<sup>b</sup> |
| KE1 | 30.25 | 24.25 | 1.24<sup>b</sup> |
| KE6 | 21 | 18.97 | 1.10<sup>a</sup> |
| Negative control | - | - | - |

Note: Unequal letter is significantly different according to Duncan test at 95% confidence level ( $P > 0,05$ )

**Figure 2.**
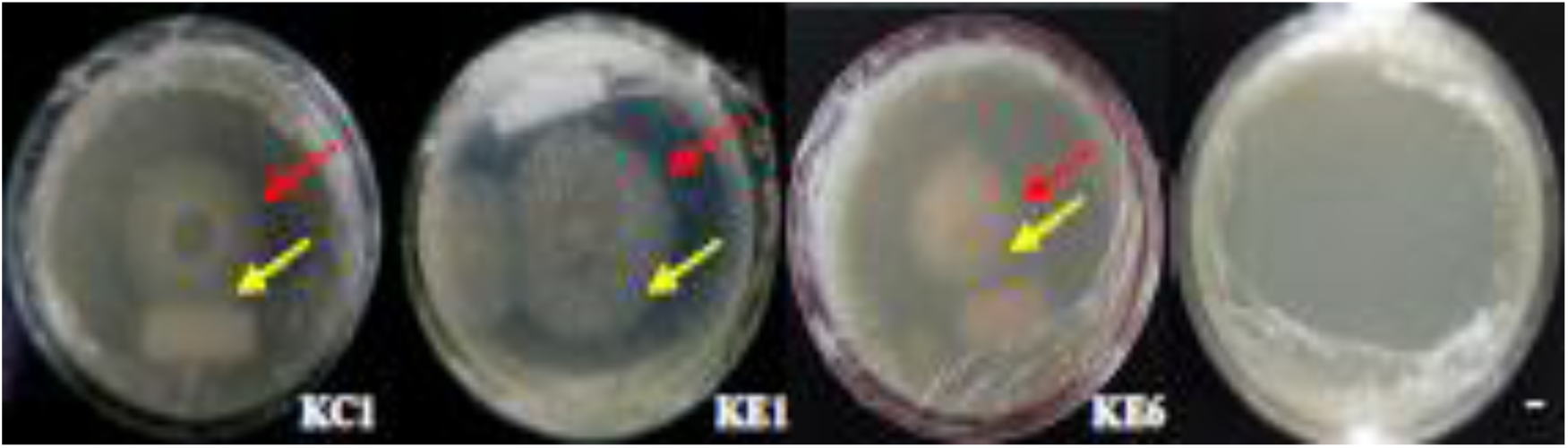
Degradation PHB of three lipolytic isolates. Note: **→** clear zone, **→** mold colony, (-) = negative control

### c. Morphology Characteristics

Three selected isolates were observed based on macroscopic and microscopic characters. Macroscopically, these colony isolates were observed in PDA medium after 7 days. The macroscopic characteristics of the colony were shown in Table 3 and Figure 3. Microscopic characteristics of each isolate were shown in Table 4 and Figure 4.

**Table 3.** Macroscopic characteristics of three selected isolates.

| Characteristics | Isolate Code |  |  |
| --- | --- | --- | --- |
|  | KC1 | KE1 | KE6 |
| Colony surface | Dark yellow | Black | Dark blue |
| Colony reverse | Yellow | Cream | Cream |
| Exudate | Absent | Absent | Present |
| Growth rate | Fast | Fast | Fast |
| Zonation | Present | Present | Absent |
| Texture | Powdery | Powdery | Powdery |
| Sporulation | Present | Present | Present |
| Diameter (mm) | 46.75 | 42.3 | 62 |
| Radial line | Present | Present | Absent |

**Table 4.**
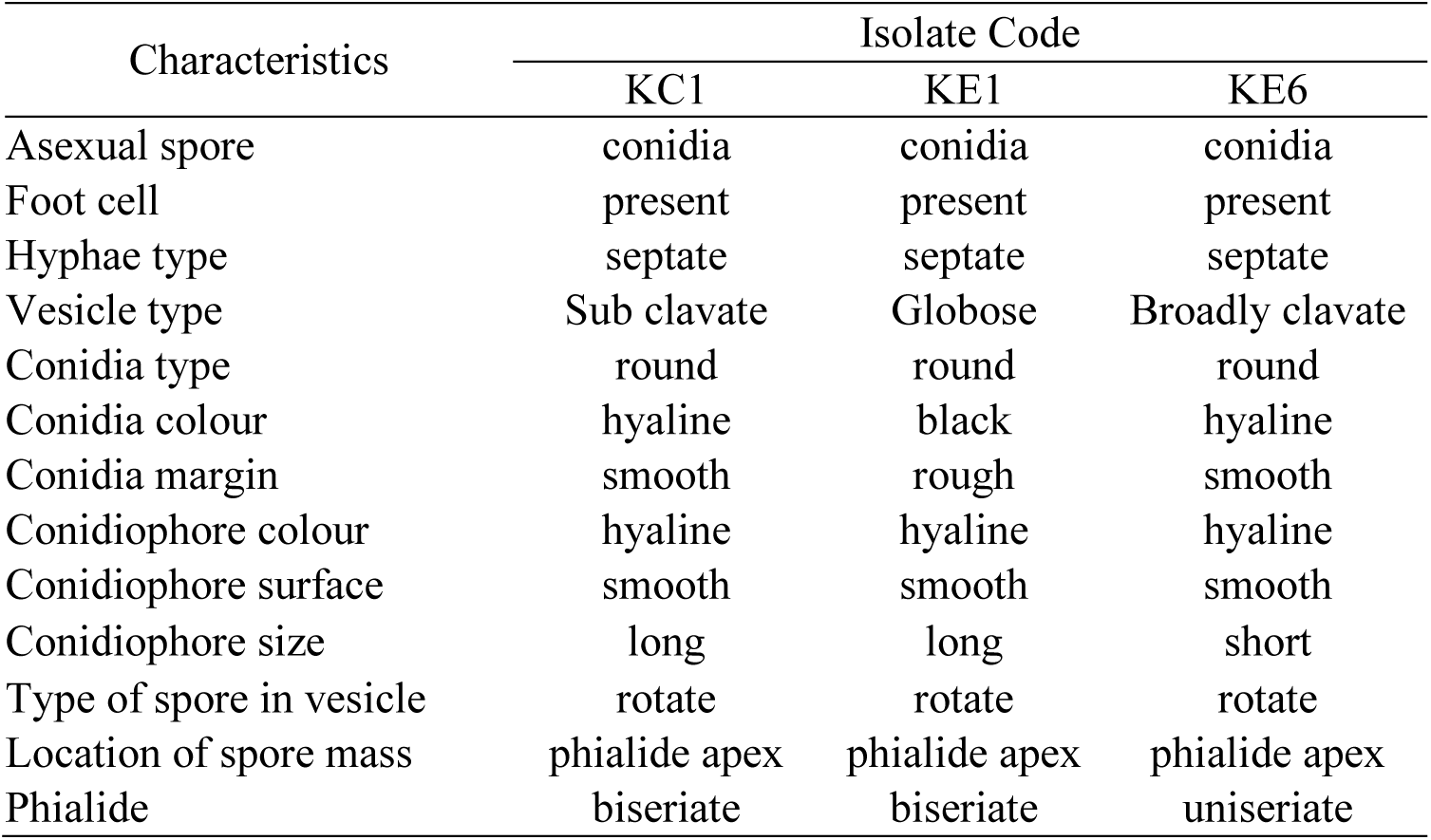
Microscopic characteristics of PHB-degrading lipolytic mold.

**Figure 3.**
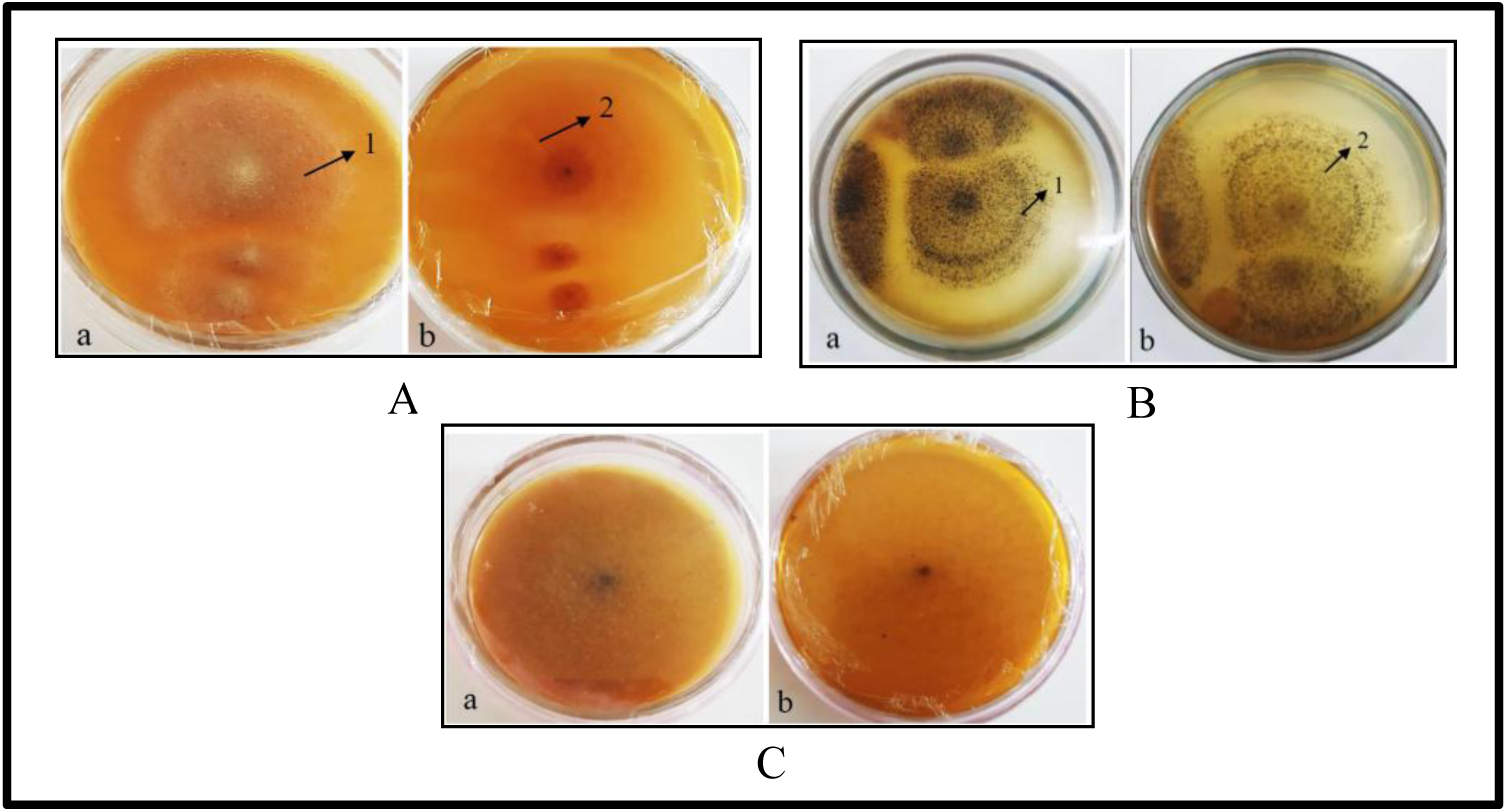
Colony morphology on media PDA at 7 days incubation. (A) isolate KC1 (B) isolate KE1 (C) isolate KE6, a: observe colony, b: reverse colony, 1: zonation, 2: radial line

**Figure 4.**
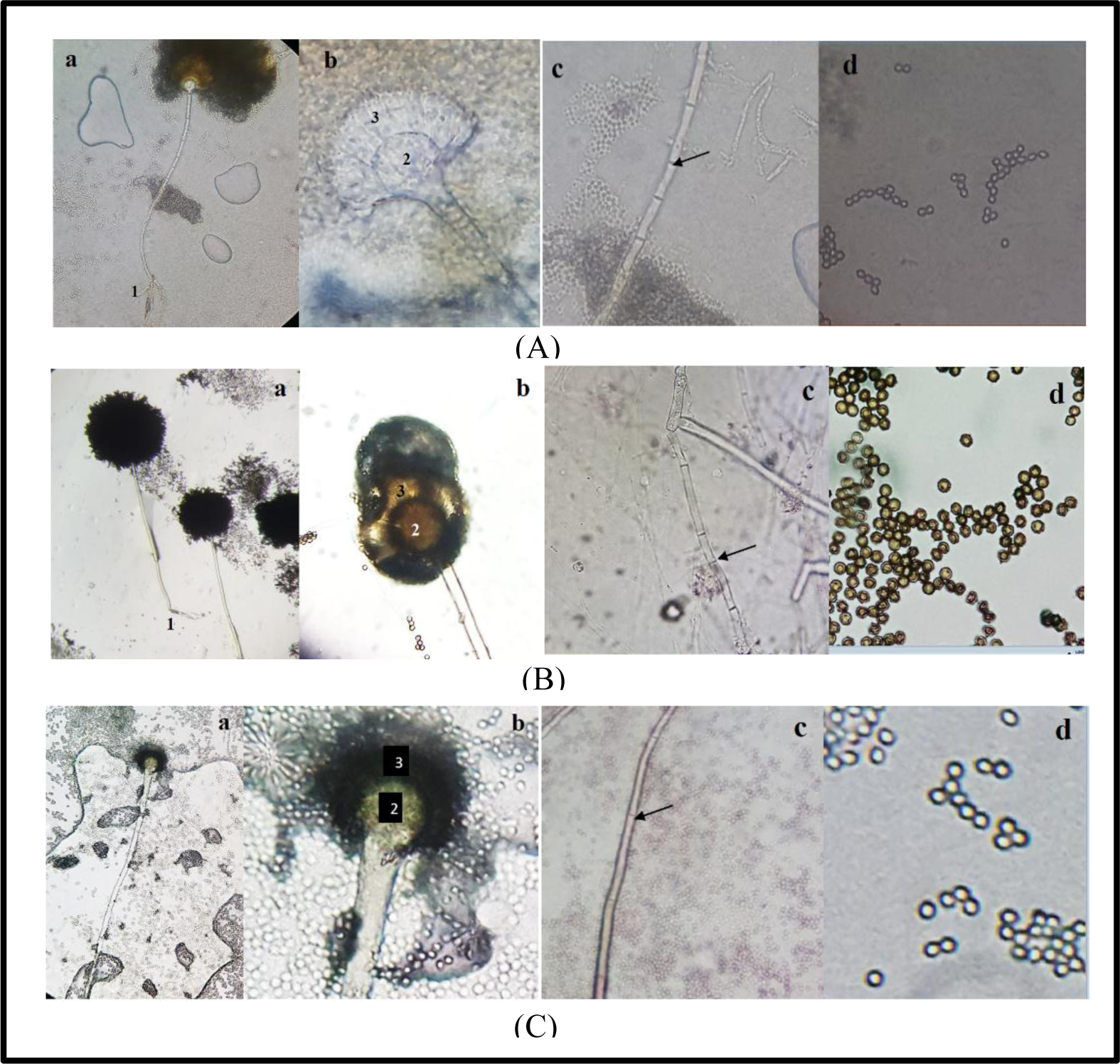
Light microscopic photographs (A) isolate KC1 (B) isolate KE1 (C) isolate KE6, 400x magnification, a: conidiophore and spore mass, 1: *foot cell* b: morphological structure, 2: vesicle, 3: phialid, c: septate hyphae, d: conidia

### d. Identity of PHB-degrading lipolytic molds

The first identification was done to determine the genera of isolates according to the phenotipic characters compared to the certain key character in *Illustrated Genera of Imperfect Fungi* (Barnett and Hunter, 1987) and *Training course for identification of Aspergillus, Penicillium and Talaromyces* (Samson, 2017). The result showed that all PHB-degrading fungi isolates were a member of *Aspergillus* due to their indentical phenotipic characters to this genera (Table 5).

**Table 5.** Profile mathcing between key characters of *Aspergillus* to isolates.

| Characters | *Aspergillus* | KC1 | KE1 | KE6 |
| --- | --- | --- | --- | --- |
| Asexual spore | Conidia | conidia | conidia | conidia |
| Footcell | Present | Present | Present | Present |
| Hyphae type | Septate | Septate | Septate | Septate |
| Conidiophore branch | Absent | Absent | Absent | Absent |
| Location of spore mass | phialide apex | phialide apex | phialide apex | phialide apex |

Numeric-phenetic identification was done based on the similarity of 33 phenotipic characters between these 3 mold isolates and 4 reference species, namely *Aspergillus niger, Aspergillus terreus, Aspergillus flavus* and *Aspergillus fumigatus*. Four reference species were selected based on the character similarity to the obtained isolates. The analysis results was visualizied on dendrogram as shown in Figure 5. Four clusters were formed and all genera in the cluster had similarity value more than 70%. Isolate KC1 and *Aspergillus terreus* were in cluster 1 with 100% similarity value. This was indicated that isolate KC1 was identic to *Aspergillus terreus*. Isolate KE1 and *Aspergillus niger* were in cluster 2 with 94.3% similarity value. This indicated that isolate KE1 was identic to *Aspergillus niger*, while isolate KE6 and *Aspergillus fumigatus* were in cluster 4 with 94.3% similarity value. This was indicated that isolate KE6 was identic to *Aspergillus fumigatus*.

**Figure 5.**
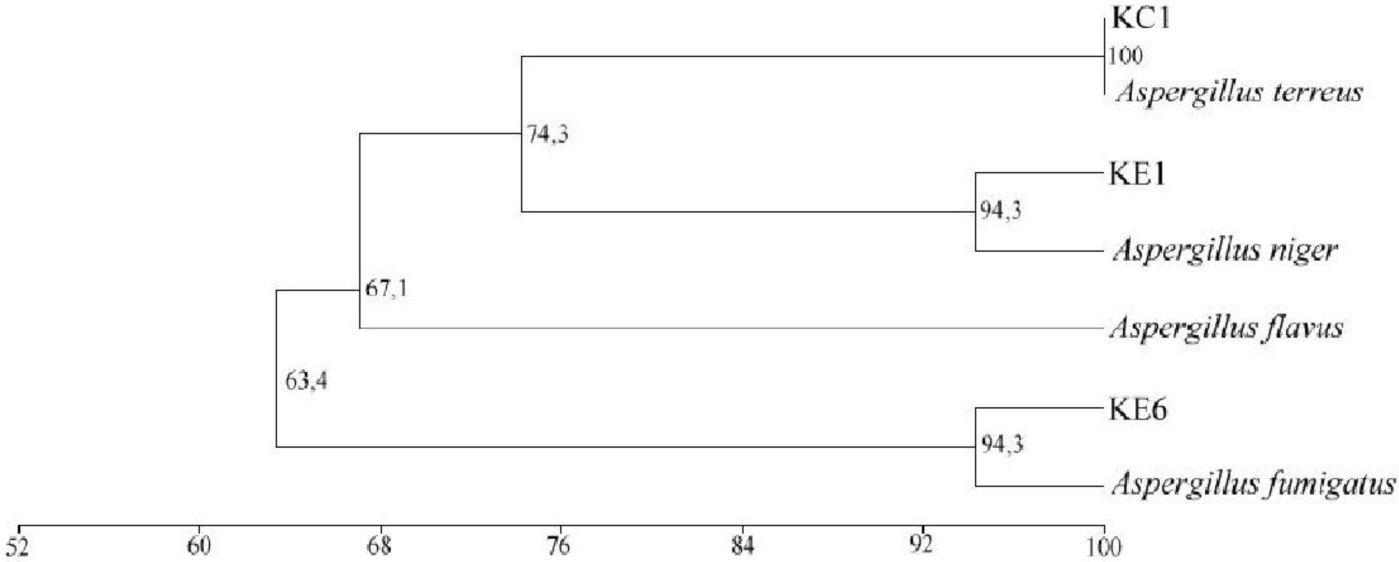
Dendrogram of similarity between 3 PHB-degrading lipolytic isolates and 4 reference spesies based on *simple matching coeficient* (SSM) and *unweighted pair-group method with arithmetic average* (UPGMA) using phenotipic characters.

## Discussion

Lipolytic molds can utilize lipid substrates as their carbon source by the action of lipase enzyme. They can be found at the appropriate environments, such as at industrial wastes, vegetable oil or milk factories, oil-contaminated soil, rotten food, compost and hot spring (Musa and Tayo, 2012; Sharma *et al*., 2001). In this study, isolate source was taken from Kendari landfill soil and there were 24 isolates had been obtained. The landfill soil is considered as a great source to obtain lipolytic molds due to many oil substrates or oil wastes are thrown there away. Some lipolytic screening methods have been proposed, one of them was rhodamine B agar method.

Rhodamine B is a fluorescence dye that will bind into the hydrolyzed fatty acid resulting the formation of orange fluorescent halos around the fungal colony (Kouker and Jaeger, 1987; Olusesan *et al*., 2009). In this medium, olive oil was used as lipid subsrate to stimulate the lipase activity. Orange fluorescent was formed due to the complexes binding between rhodamine B cation and uranil ion of the hydrolyzed fatty acid. According to the result, sixteen lipolytic mold isolates were found. Out of them, there were three lipolytic isolates with the highest lipolytic activity. Some studies have been also reported that this method was used to screen both fungal and bacterial lipolytic activities (Mendes *et al*., 2019; Ramnath et al., 2017). Furthermore, Kouker and Jaeger (1987) stated that this method was preferrable than others due to there was not inhibition effect to the growth or change in the physiological characters of microorganisms.

The ability of isolates to degrade biofilm PHB was observed based on the formation clear zone around the colony. The fungal growth and clear zone were checked everday. After 2 days, the clear zone was formed. This result was contrary with the study by Lee *et al*. (2005) who proposed that the clear zone observed after 1-3 weeks. As shown in Table 2, three lipolytic isolates were potential to degrade biofilm PHB. Jackson (1998) stated that the clear zone formation rate or PHB degradation rate was affected by four factors, such as growth rate, enzyme secretion, enzyme activity, and enzyme diffusion into agar medium. Suriawiria and Unus (1986) also stated that agar concentrations and growth conditions were affected the degradation rate of PHB. The degradation rate will be inhibited when agar concentration is exceeding 3% (w/v) may likely due to the inhibition of enzyme diffusion into agar medium, while the growth condition affects fungal growth as well as degradation rate.

Enzyme which can catalyze the degradation of PHB is PHB depolymerase, but lipases also have been reported to catalyze this (Leja and Lewandowicz, 2010). Lipases catalyze the hydrolysis of ester bond at oil-water interface. Lipases can degrade PHB due to the present of ester bond in PHB monomer (Leja and Lewandowicz, 2010). Poirier *et al*. (1995) was proposed two process of PHB degradation, first is the attachment of enzyme into the particle surface and will form enyme-substrate complexes and the second is hydrolysis of PHB into soluble oligomer or monomer. [R]-3-hydroxybutirate acid is formed as the last product of this ennzymatic activity.

There are three lipolytic molds obtained from Kendari landfill soil which were potential to degrade PHB, namely isolate KC1, KE1 and KE6. Based on numeric-phenetic indentification, isolate KC1 is a member of *Aspergillus terreus*, isolate KE1 is a member of *Aspergillus niger* and isolate KE6 is a member of *Aspergillus fumigatus. Aspergillus* has been known as one of the most predominant genera of lipase-producing and PHB-degrading molds (Matavulj and Molitoris, 1992). Some studies also reported that these molds had the ability to degrade PHB (Kumaravel *et al*., 2010; Lodhi, *et al*., 2011; Merugu, 2012; Aburas, 2016; Aly et al., 2017).

## Conclusion

Landfill soil can be a source of PHB-degrading lipolytic molds. *Aspergillus terreus* KC1, *Aspergillus niger* KE1 and *Aspergillus fumigatus* KE6 are lipolytic molds which have the ability to degrade PHB biofilm. These lipolytic molds might be potential in degrading another bioplastics. Furthermore, their potential can be enhanced through optimization study so it might be applied at industrial scale.

